# Bacterial Signatures and Community Structure in the Phyllosphere of *Eugenia uniflora*: Developmental Dynamics and Core Microbiome in Myrtaceae

**DOI:** 10.64898/2026.04.10.717871

**Authors:** Isabel Cristina Cadavid Sánchez, Dámaris Esquén, Rogerio Margis, Frank Guzman

## Abstract

Plants recruit microorganisms to form mutually beneficial associations that enhance their health, productivity, and resilience. The composition of the plant microbiome is shaped by factors such as host species, developmental stage, genotype, and tissue type, with microbial recruitment mediated by plant exudates and secondary metabolites. *Eugenia uniflora*, a Myrtaceae species native to Brazil’s Atlantic Forest, produces pharmacologically relevant secondary metabolites and holds ecological and economic value. However, little is known about its associated microbiome, particularly from a metagenomic perspective. In this study, we investigated the phyllosphere bacterial communities, both epiphytic and endophytic, of *E. uniflora* across two developmental stages (young and mature trees). We also examined the core microbiome shared between *E. uniflora* and other Myrtaceae genera to better understand microbial diversity and structure within this family. Amplicon sequencing of the V3–V4 region of the 16S rRNA gene was conducted on 19 *E. uniflora* samples and 13 additional samples from three other Myrtaceae genera. In *E. uniflora*, we identified 1,456 bacterial ASVs representing 17 phyla, 115 families, and 171 genera. Alpha and beta diversity analyses revealed significant differences in bacterial community composition between developmental stages. Genera such as *Massilia* and *Hymenobacter* were more abundant in mature trees, while *Aureimonas* and *Terriglobus* were more common in young plants. Leaf microbiome functional potential shifted with plant age, with older leaves favoring secondary metabolite production and younger leaves emphasizing microbial interactions and defense. A total of 16 genera formed the Myrtaceae core microbiome, with five, *Methylobacterium–Methylorubrum*, *Hymenobacter*, *Sphingomonas*, *Bdellovibrio*, and *Terriglobus*, present in 100% of samples. Notably, ∼0.7% of the bacterial diversity remained poorly classified, highlighting the underexplored nature of Myrtaceae-associated microbiomes and their potential for bioprospecting.

## 1. Introduction

The plant microbiome encompasses the community of microorganisms that colonize internal and external plant compartments, playing vital roles in plant health and development^1,2^. These microbial communities support nutrient acquisition through processes such as nitrogen fixation and mineral solubilization, promote growth via phytohormone production, and enhance tolerance to biotic and abiotic stressors by activating plant defense mechanisms and suppressing pathogens ^3–5^. Due to these benefits, plant-associated microbes are increasingly explored for the development of bioinoculants and sustainable agricultural technologies ^6^.

The structure and diversity of the plant microbiome are influenced by several factors, including host species, genotype, developmental stage, and tissue type, and may reflect long-term co-evolutionary relationships between plants and their microbial partners ^7,8^. Changes in plant diversity—due to domestication or environmental disturbances—can negatively affect the associated microbiota ^9,10^, making the plant microbiome a valuable indicator of ecosystem health and a potential reservoir for biotechnological applications.

Within this context, plants of the Myrtaceae family, native to the Neotropics and Australia, are of ecological and economic significance. In Brazil, this family is abundant in biodiverse ecosystems such as the Atlantic Forest, contributing as a food source for insects, birds, and mammals through their flowers and fruits. Many Myrtaceae species bear edible fruits used in human diets and possess medicinal properties due to their production of bioactive secondary metabolites, including flavonoids and terpenoids.

*Eugenia uniflora* (commonly known as pitanga), a native Myrtaceae species from the Brazilian Atlantic Forest, exemplifies this dual ecological and pharmacological relevance. Like other Myrtaceae relatives, it produces edible fruits and exhibits potential for pharmaceutical applications. However, despite its significance, the microbiome associated with *E. uniflora* remains poorly characterized, especially through high-throughput sequencing approaches ^11–13^. Understanding the bacterial communities associated with *E. uniflora* could reveal novel taxa with potential for bioprospecting, sustainable agriculture, and ecosystem monitoring. In this study, we investigated the phyllosphere bacterial communities, including both epiphytic and endophytic compartments, of *E. uniflora* across two developmental stages (young and mature trees). Additionally, we characterized the core microbiome shared between *E. uniflora* and other Myrtaceae genera to assess microbial diversity and structure within this plant group. Our aim was to identify bacterial taxa that are consistently present across Myrtaceae species, as well as those that may be specific to individual genera. These insights will help establish foundational knowledge of plant–microbe interactions within Myrtaceae and highlight microbial candidates for biotechnological applications such as biofertilizers and biocontrol agents, contributing to the development of more sustainable agricultural practices.

## 2. Methodology

### 2.1 Plant sample collection

To study the bacterial communities associated with *Eugenia uniflora*, 19 leaf samples were collected from apparently healthy plants, including 8 from young individuals and 11 from mature trees. Of these, 13 samples were obtained from Margis Farm in the district of Morungava, municipality of Gravataí, and 6 from the Campus do Vale of the Federal University of Rio Grande do Sul (UFRGS) in Porto Alegre. Both locations are situated in the state of Rio Grande do Sul, Brazil.

To compare the phyllosphere microbiomes of pitanga and related Myrtaceae trees, leaf samples were collected from 13 individual trees representing four genera at Margis Farm. Between one and four replicates per genus were sampled. For each tree, leaves were collected from multiple locations across the canopy to capture within-plant variation in microbial communities. In total, eight Myrtaceae species were included in the analysis (Supplementary Table S1): *Eugenia brasiliensis* (grumixama), *Eugenia pyriformis* (uvaia), *Eugenia selloi* (pitangão), *Myrcianthes pungens* (guabijú), *Plinia trunciflora* (jaboticaba), *Plinia edulis* (cambucá), *Psidium guajava* (guava), *Psidium cattleianum* (araçá), and the sampled trees were located within a one-hectare area, with a minimum spacing of 50 cm between individuals (Fig S1). All samples were collected in November 2021, during the spring season.

### 2.2 Collection of microbiome community

Leaf material from each sample was transported to the laboratory in sterile plastic bags. To isolate the total phyllosphere microbiome, leaves were cut into small pieces (∼0.5 × 0.5 mm) using sterile tweezers and a scalpel on a sterilized glass Petri dish under a laminar flow hood. The leaf fragments were transferred to an Erlenmeyer flask containing 50 mL of sterile, pre-chilled PBST buffer ^14^. Samples were shaken at 180 rpm for 2 hours at 4 °C. After incubation, 40 mL of the suspension was transferred to sterile Falcon tubes and centrifuged at 3,000 rpm for 30 minutes. The resulting pellets were stored at −20 °C until DNA extraction.

### 2.3 DNA isolation and sequencing

Total DNA was extracted using the ZymoBIOMICS™ DNA MiniPrep Kit (Zymo Research Corporation). DNA quality and concentration were assessed using agarose gel electrophoresis and a NanoDrop spectrophotometer. The V3–V4 region of the 16S rRNA gene was amplified using the primer pair 341F (5′-CCTAYGGGRBGCASCAG-3′) and 806R (5′-GGACTACNNGGGTATCTAAT-3′) ^15,16^. PCR amplification was performed following the protocol described by Christoff et al. (2017)^17^. The PCR products were visualized on a 1.5% agarose gel, and samples showing a clear band between 400 and 450 bp were selected for sequencing. Library preparation and sequencing were carried out on the Illumina MiSeq platform using 1 × 300 bp single-end reads.

### 2.4 Sequence analysis

Raw sequence data were first assessed for quality using FastQC, and low-quality reads (Phred score < 30) and sequences shorter than 200 nucleotides were filtered out using Trim Galore ^18^. To remove host-derived contamination, Bowtie2 was used to map and filter reads corresponding to chloroplast (cp) and mitochondrial (mt) sequences by aligning against *Eugenia uniflora* cp16S rRNA and *Eucalyptus grandis* mt18S rRNA gene references. These organellar reads were excluded from further analysis.

Subsequent quality control and processing were performed using the DADA2 pipeline ^19^, which included read filtering, trimming, denoising, and chimera removal. Amplicon sequence variants (ASVs) were inferred at 100% sequence similarity. Representative sequences for each ASV were aligned using MAFFT in QIIME2, and phylogenetic trees (unrooted and rooted) were generated with FastTree to support downstream diversity analyses. Taxonomic classification of ASVs was performed in QIIME2 ^15^ using a pre-trained Naïve Bayes classifier against the SILVA 16S rRNA database ^20,21^, applying a minimum confidence threshold of 0.8. An ASV table was constructed to represent the distribution of bacterial taxa across samples, and taxonomic bar plots were generated using the *ggplot2* package in R at the phylum, family, and genus levels.

ASVs identified as plant-derived—classified as Cyanobacteria/Chloroplast or Proteobacteria/Mitochondria, were filtered out using QIIME2’s taxonomy-based feature table filtering option. In total, 32 libraries representing four plant genera were analyzed. After all filtering steps, sequencing depth ranged from 5,530 reads (*Plinia edulis*) to 30,573 reads (*Eugenia uniflora*) per sample. Detailed information on read counts and ASV numbers per sample is provided in Supplementary Tables S1 and S2.

### 2.5 Statistical analyses

The relative abundance of ASV was calculated after rarefying all samples to the minimum sequencing depth observed per sample. Subsequently, various diversity indices and statistical analyses were performed using the q2-diversity plugin in QIIME2.

Alpha diversity metrics, including *Observed Features*, *Chao1*, *ACE*, *Shannon*, and *Simpson* indices, were computed to assess species richness and diversity within samples. Beta diversity was evaluated using Bray-Curtis dissimilarity, and Principal Coordinates Analysis (PCoA) plots were generated using R packages.

To assess statistical significance, pairwise Kruskal–Wallis tests were used to compare alpha diversity across different ages (young and old trees) of *Eugenia uniflora*, as well as across species and genera in the Myrtaceae dataset. For beta diversity, Bray-Curtis distance matrices were analyzed using permutational multivariate analysis of variance (PERMANOVA) with 999 permutations via the *vegan::adonis2* function, complemented by the Wilcoxon test for pairwise comparisons.

Taxonomic assignments were collapsed to the genus level (sixth taxonomic rank) for downstream analyses including core microbiome identification, Venn diagram visualization, and differential abundance testing with EdgeR.

The core microbiome of *Eugenia uniflora* and the Myrtaceae family was defined by plotting the log10 mean relative abundance against occupancy for each bacterial genus in R, following the approach of Shade and Stopnisek (2019)^22^. Genera present in at least 75% of samples (>75% occupancy) were considered part of the highly conservative core microbiome.

Additionally, Venn diagrams were generated to illustrate shared and unique bacterial genera among the plants, using the online tool InteractiVenn (http://www.interactivenn.net/).

Venn diagrams were constructed considering the presence of bacteria genus in at least one sample per plant group.

Microbial differential abundance analysis was performed using the *edgeR* package in R to identify bacterial genera with differential representation between developmental stages (old vs. young) of *Eugenia uniflora* and across Myrtaceae groups, as well as. Raw count data were filtered to remove low-abundance taxa and normalized using the trimmed mean of M-values (TMM) method. Negative binomial generalized linear models were fitted with group and developmental stage as main factors. Differentially abundant genera were identified based on log fold change (logFC) and false discovery rate (FDR) adjusted *p-values*.

### 2.6 Functional prediction of bacterial communities

Functional profiles of bacterial communities were predicted using the *Tax4Fun R* package, which infers the metabolic potential of microbial communities based on 16S rRNA gene sequences. The pipeline relies on SILVA-based taxonomic classifications to map taxa to known KEGG Orthologs (KOs), generating a table of predicted gene abundances. The resulting KO abundance table was used to generate box plots of the top functional categories across samples using the ggplot2 package in R.

Additionally, ecological functional profiling was also performed using the *FAPROTAX* database. The ASV table obtained from 16S rRNA sequencing was annotated using the collapse_table.py script, with taxonomy-based mapping to ecological functions. The resulting functional table was used to generate a bar plot of the top functional categories across samples using the ggplot2 package in R.

Differences in metabolic functions between sample groups were assessed using abundance matrices of metabolic pathways or ecological functions. Bray-Curtis distances were calculated using the **vegan** package in R to quantify dissimilarities among samples. These distances were visualized using non-metric multidimensional scaling (NMDS, two axes), with confidence ellipses representing the variability within each group. Statistical differences between groups were tested using PERMANOVA (**adonis2**, vegan) based on the distance matrix. To identify functions that were differentially abundant between groups, log fold change (logFC) and false discovery rate (FDR)-adjusted p-values were calculated using **edgeR**. For visualization of functional patterns, only the most abundant functions (top 10 KOs by total abundance across all samples) were retained.

## 3. Results

### 3.1 Bacterial community structure and functional potential vary with developmental stage in Eugenia uniflora

The bacterial community associated with *Eugenia uniflora* comprised 1,458 ASVs (Amplicon Sequence Variants, Table S1), which were used to assess microbial diversity. Alpha diversity analyses indicated significantly higher diversity in older trees compared to younger ones, with increases in both species’ richness and evenness. This was supported by significantly higher values in both the Shannon and Simpson diversity indices (*p* < 0.01, *Kruskal–Wallis* test, Table S3, Fig 1).

**Fig 1.**
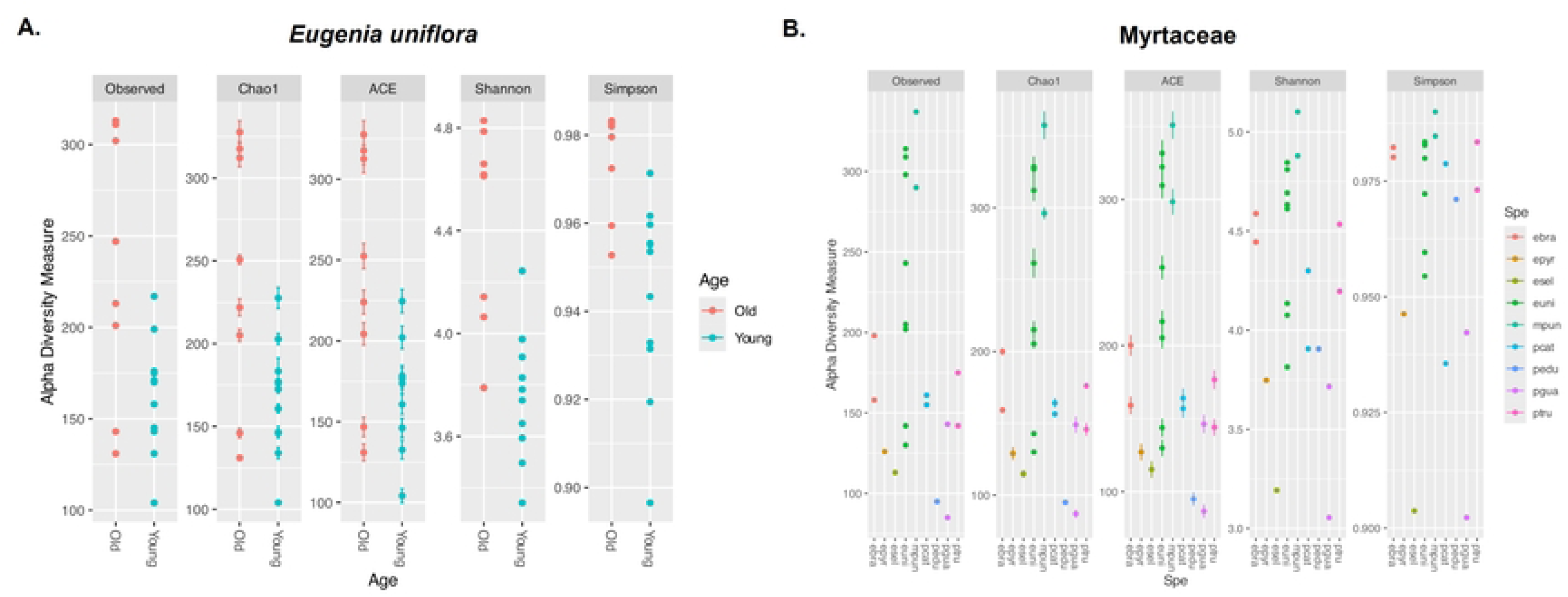
Alpha diversity of phyllosphere bacterial communities based on different diversity indices. (a) *Eugenia uniflora* at two developmental stages (young and mature trees). (b) *Myrcianthes* genus samples. Statistical differences between groups were evaluated using the Kruskal–Wallis test, and p-values < 0.05 were considered statistically significant.

Beta diversity analysis revealed a clear pattern in community composition, with bacterial communities clustering according to developmental stage (Fig 2A). This pattern was confirmed by PERMANOVA (*p* < 0.01, Adonis test) based on Bray–Curtis dissimilarity (Table S3). These results indicate that the microbial composition of older trees is distinct from that of younger ones, a pattern that is visually evident in the PCoA plot, where axis 1 explains 33.5% of the variation.

**Fig 2.**
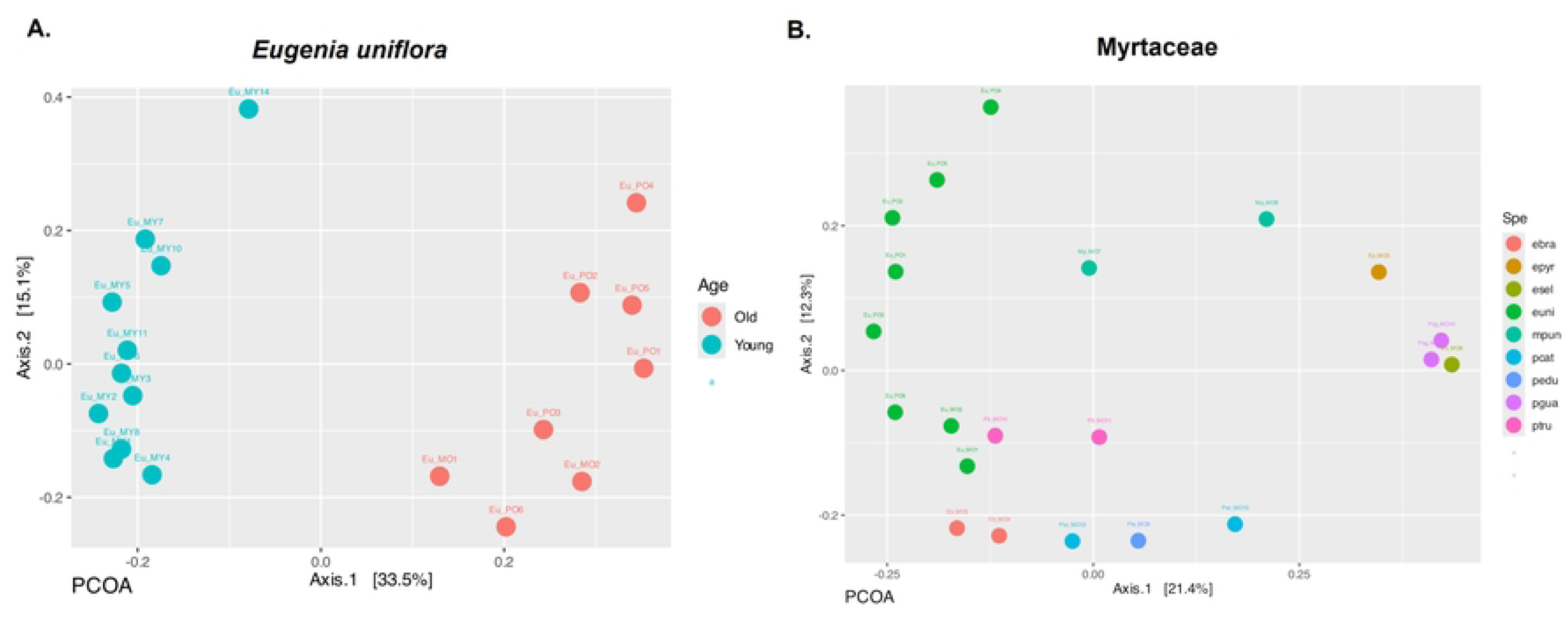
Principal Coordinate Analysis (PCoA) based on Bray–Curtis dissimilarity of phyllosphere bacterial communities. (a) *Eugenia uniflora* at two developmental stages (young and mature trees). (b) *Myrcianthes* genus samples. Each point represents the bacterial community of an individual sample, and colors indicate different plant ages or species.

Following taxonomic classification, the bacterial profiles across *Eugenia uniflora* (pitanga) samples were examined. At the phylum level, the most dominant groups identified were Proteobacteria (77%), Bacteroidota (15%), and Actinobacteriota (3%) (Fig S1a). At the family level, *Beijerinckiaceae* (54%), *Hymenobacteraceae* (12%), and *Bdellovibrionaceae* (2%) were predominant (Fig S1b). At the genus level, the most abundant taxa included *Methylobacterium-Methylorubrum* (33%), *Methylocella* (16%), *Hymenobacter* (15%), 1174-901-12 (12%), and *Sphingomonas* (9%) (Fig 3a, Table S4).

**Fig 3.**
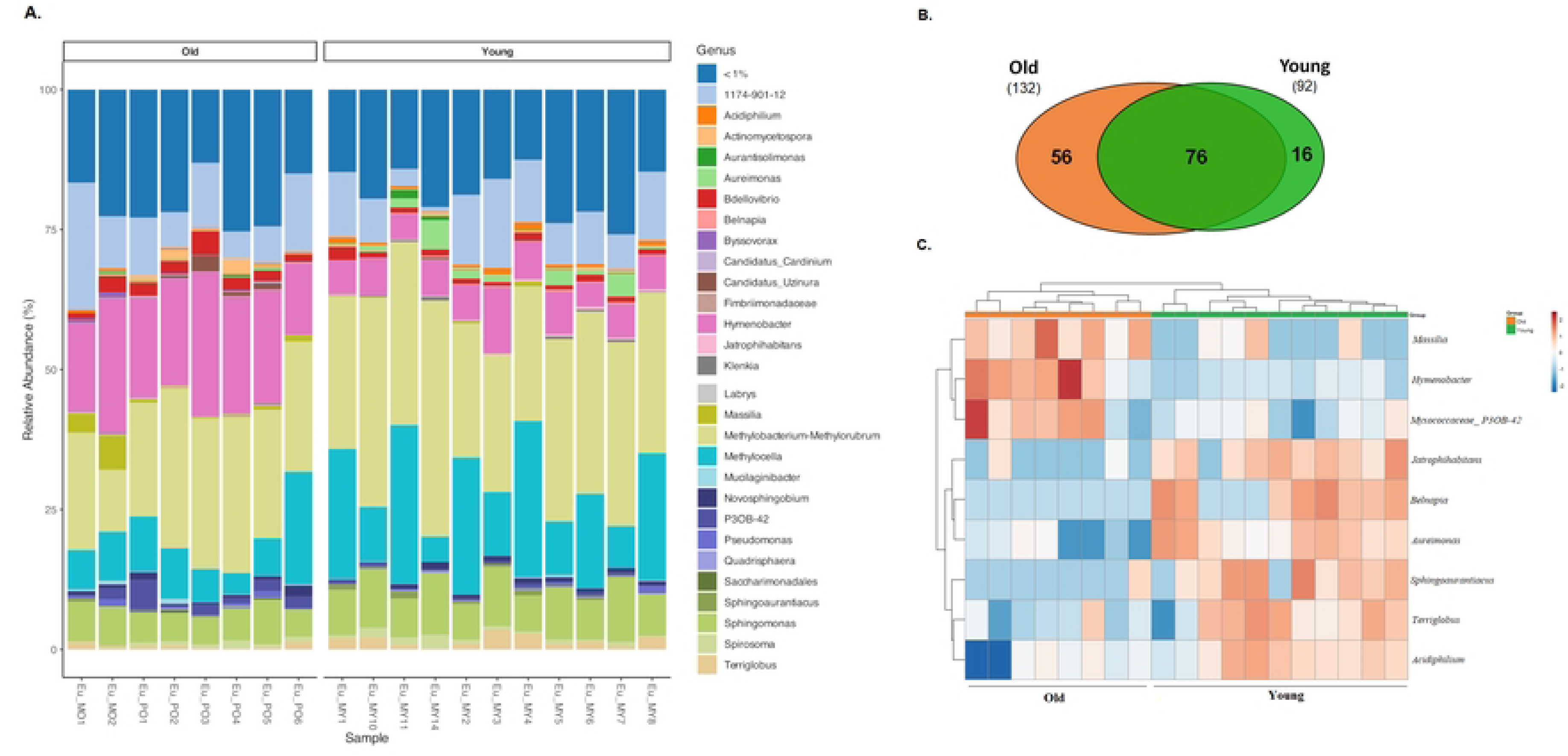
Taxonomic composition and differential abundance of the *Eugenia uniflora* phyllosphere bacterial community at two developmental stages. (a) Taxonomic bar plots showing the relative abundance of bacterial genera. (b) Venn diagram displaying shared and unique genera between young and mature trees. (c) Heatmap of differentially abundant bacterial genera; blue indicates lower and pink higher relative abundance.

A Venn diagram (Fig 3b) showed that while young and old trees shared many bacterial genera, certain taxa were unique to each developmental stage (Table S5). To move beyond presence/absence data and consider relative abundance, a differential abundance analysis was conducted using edgeR. This analysis revealed that specific bacterial genera were significantly associated with tree age. For example, *Massilia*, *Hymenobacter*, and a member of the *Myxococcaceae* family (P3OB-42) were more abundant in mature trees, whereas *Jatrophihabitans*, *Belnapia*, *Aureimonas*, *Sphingoaurantiacus*, *Terriglobus*, and *Acidiphilum* were more prevalent in younger plants (Fig 3c, Table S6).

To investigate the metabolic and ecological functions of the leaf microbiome, predictions were made based on 16S rRNA sequences. Functional profiling revealed that the developmental stage of *Eugenia uniflora* shaped the bacterial community, as shown by NMDS based on Bray–Curtis dissimilarity (*p* < 0.01, Adonis test) of KEGG Orthology (KO) categories (Fig 4a, Table S7). *EdgeR* statistical analysis showed functions were differentially abundant between groups (Table S7). Older plants were enriched in pathways related to terpenoid-quinone (ko00130), antibiotic (ko00253, ko00311), glycerolipid (ko00561), retinol (ko00830), and polyketide (ko01056) biosynthesis, as well as bacterial invasion of epithelial cells (ko05100), whereas younger plants showed higher abundance of sphingolipid signaling (ko04071) and infectious disease–associated pathways (ko05120) (Fig 4b). These results indicate that leaf microbiome functional potential shifts with plant age, with older leaves favoring secondary metabolite production and younger leaves emphasizing microbial interactions and defense. Predicted functions across all samples included methanotrophy, methylotrophy, nitrogen fixation, aerobic chemoheterotrophy, hydrocarbon degradation, predatory or exoparasitic behavior, and general chemoheterotrophy (Fig 4c; Table S8). DESeq2 analysis (Table S8) revealed that predatory or exoparasitic behavior and intracellular parasitism were significantly enriched in older plants, suggesting more complex microbial interactions in mature trees that may influence community structure and ecological dynamics.

**Fig 4.**
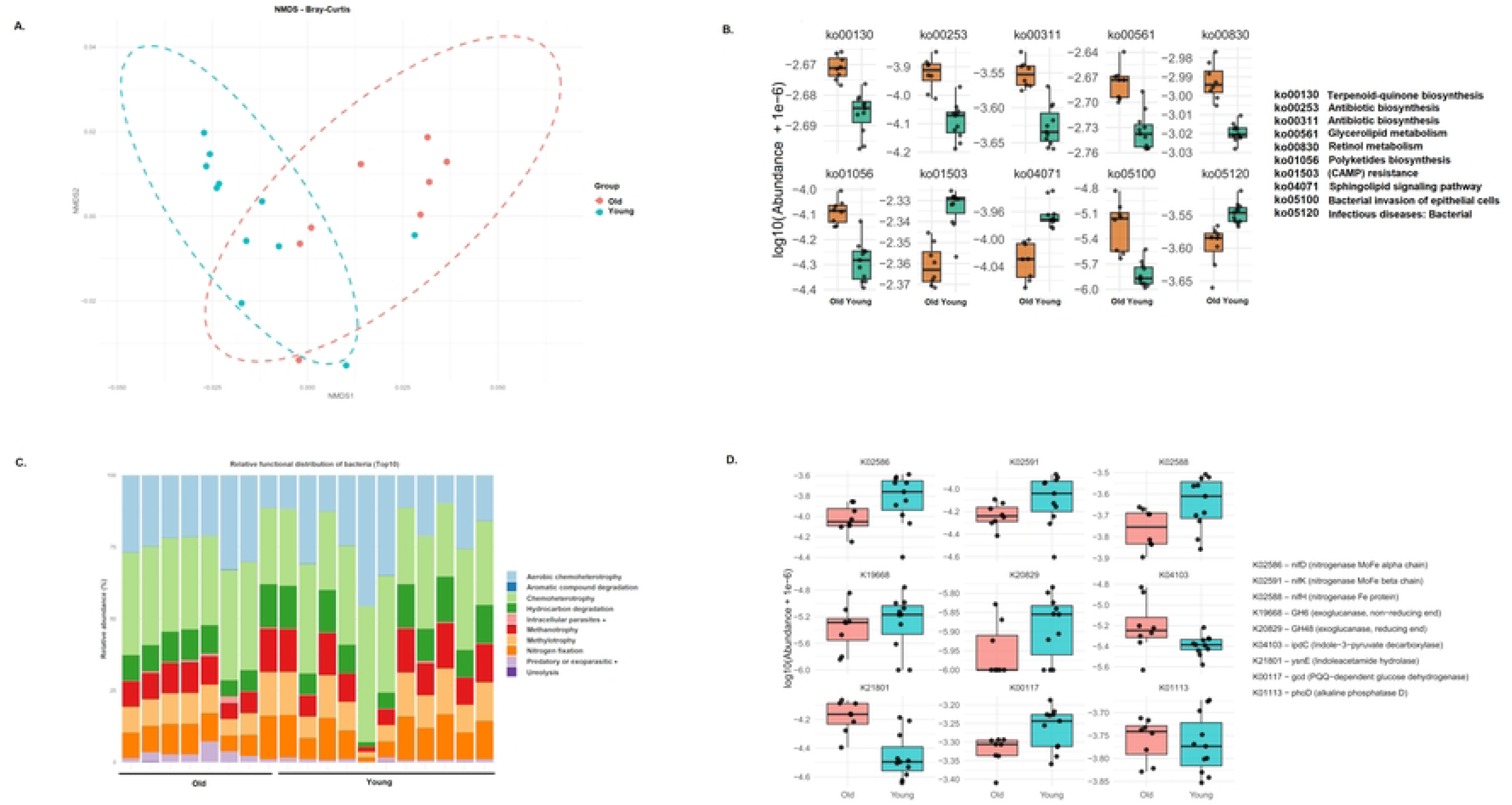
Functional analysis of the *Eugenia uniflora* phyllosphere bacterial community at two developmental stages. (a) Non-metric multidimensional scaling (NMDS) based on Bray–Curtis distances of KEGG pathway abundance matrices; each point represents a bacterial community sample. (b) Box plots showing KEGG pathways with differential abundance between young and mature trees. (c) Functional profiles based on ecological function predictions; asterisks indicate functions with statistically significant differences between stages. (d) Box plots displaying the abundance of selected PGPR functional genes inferred from rhizosphere communities of young and mature trees.

Likewise, PGPR-associated functional genes also varied according to developmental stage (Fig 4d). Genes encoding the nitrogenase complex—nifH (K02588), nifD (K02586), and nifK (K02591)—showed low but consistent abundance, with a trend toward higher levels in the phyllosphere of young plants. Auxin biosynthesis genes ipdC (K04103) and ysnE (K21801) were detected in both groups, with ipdC showing relatively greater abundance in mature samples. Cellulolytic genes GH6 (K19668) and GH48 (K20829), as well as phosphate-solubilizing markers gcd (K00117) and phoD (K01113), were broadly distributed, though gcd was more abundant in younger phyllospheres. In contrast, siderophore biosynthesis genes entA (K00216) and pvdA (K10531) tended to be enriched in older plants. These patterns suggest that the functional potential of the phyllosphere microbiome shifts with host age, modulating nutrient acquisition and growth-promotion traits throughout development.

### 3.2 Conserved microbial diversity and core bacterial taxa in Eugenia uniflora and other Myrtaceae

The bacterial community associated with Myrtaceae genera comprised 2,148 ASVs (Amplicon Sequence Variants, Table S2). Among the species evaluated, considering only mature trees, alpha diversity metrics showed no significant differences (Fig 1B, Table S9). Similarly, beta diversity analysis revealed no significant differences in microbial community composition between *Eugenia uniflora* and the other Myrtaceae genera. Pairwise comparisons supported this pattern, showing no statistically significant differences between *Eugenia* and the other three genera (Fig 1B, Table S9).

Taxonomic profiling revealed that the dominant bacterial phyla across all samples were Proteobacteria (73%), Bacteroidota (19%), and Actinobacteriota (3%) (Fig S2a). At the family level, the most abundant groups were *Beijerinckiaceae* (42%), *Sphingomonadaceae* (23%), and *Hymenobacteraceae* (17%) (Fig S2b). As expected, *Eugenia uniflora* shares a core microbiome with other members of the Myrtaceae family (Table 1), suggesting a conserved set of microbial taxa that may play essential roles in the health and functioning of these plants.

**Table 1.**
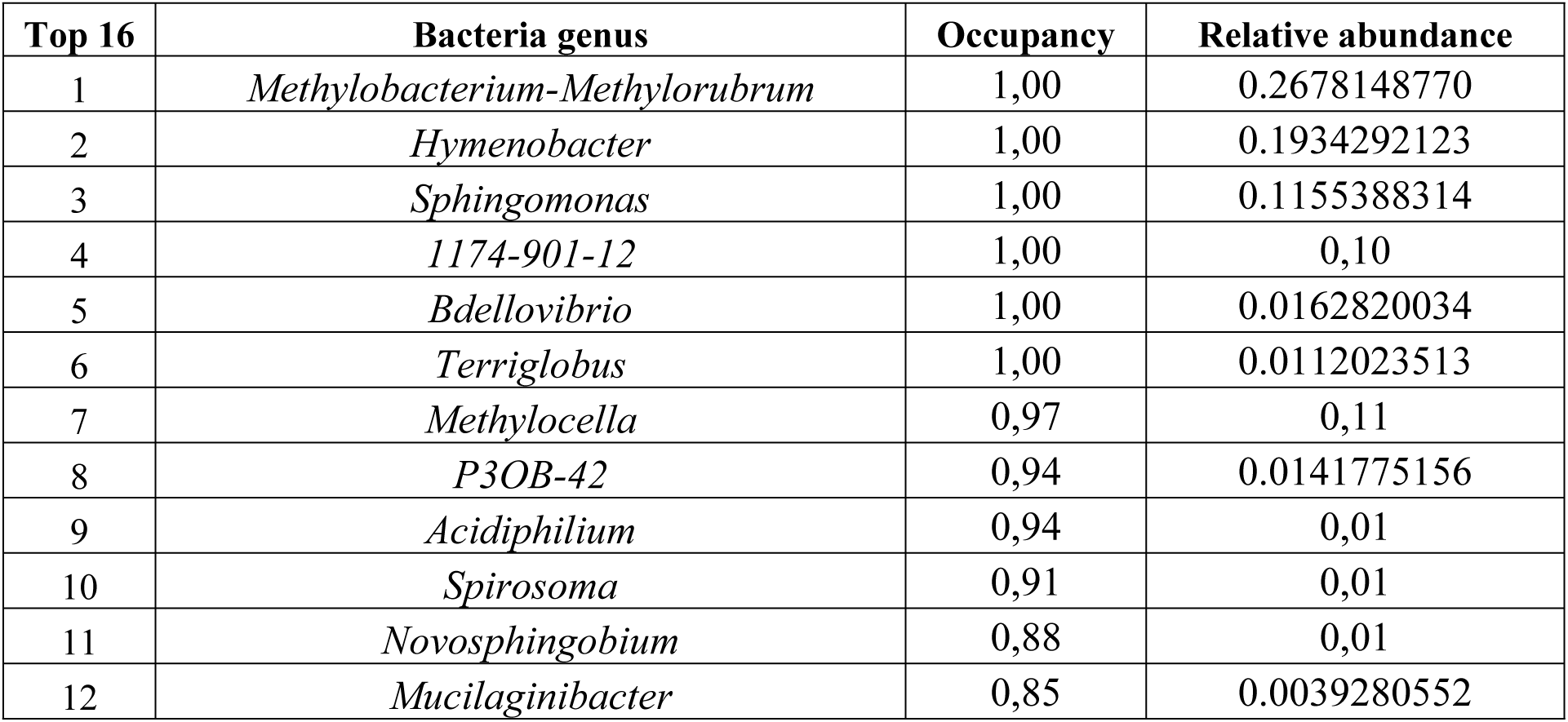
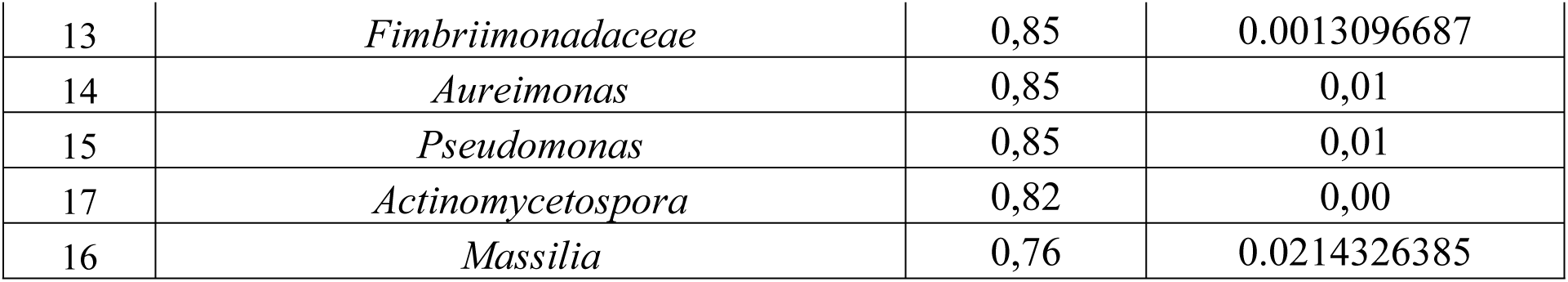
Top sixteen core microbiome members of *Eugenia uniflora* and other Myrtaceae plants.

A total of 21 bacterial genera were identified as part of the core microbiome of *E. uniflora*, defined as genera present in more than 75% of samples and consistently abundant across individuals (Fig 5A, Table S10). Comparative analysis with other Myrtaceae species revealed shared genera, suggesting a conserved core microbiome within the family (Fig 5B, Table S11).

**Fig 5.**
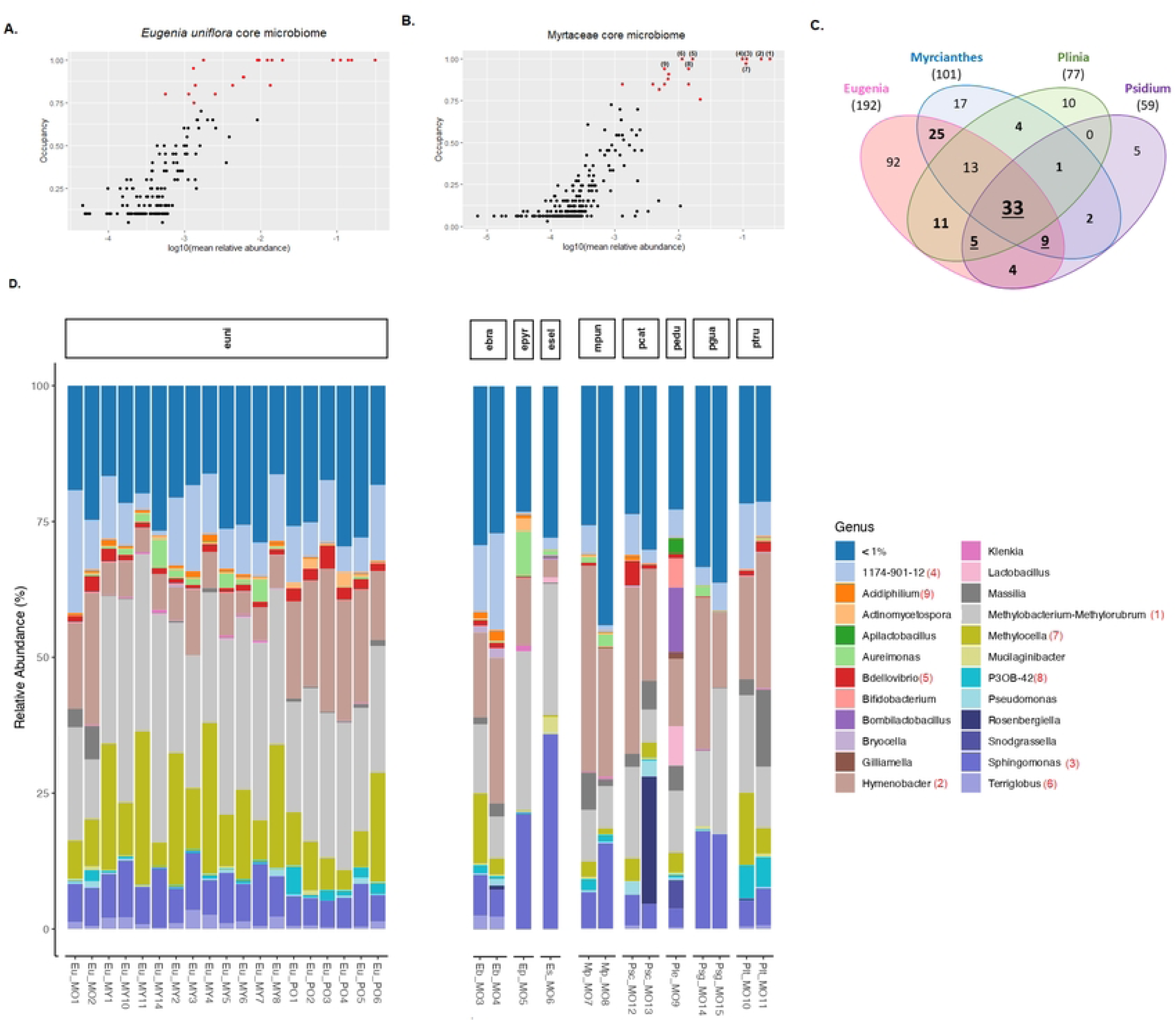
Core microbiome of *Eugenia uniflora* and other Myrtaceae species. (a) *E. uniflora* core microbiome showing bacterial occupancy (genus level) and log (mean relative abundance). Genera occurring in more than 75% of plant samples were considered part of the core microbiome and are shown as red points. (b) *Myrtaceae* core microbiome with bacterial occupancy and relative abundance at the genus level; core taxa (present in >75% of samples) are indicated in red, and the top nine genera are highlighted. (c) Venn diagrams showing the number of unique and shared bacterial genera among plant groups; presence in at least one sample per group was considered for inclusion. (d) Taxa bar plots showing the most abundant bacterial genera; taxa with high occupancy (top nine) are highlighted.

To assess the overlap of bacterial taxa among host species, a Venn diagram was generated (Fig 5C), showing that 33 genera were present across all Myrtaceae hosts analyzed. Of these, 17 genera were identified as part of the broader Myrtaceae core microbiome, meeting the criteria of being present in at least 75% of samples and exhibiting relatively high abundance. Notably, five genera, *Methylobacterium–Methylorubrum*, *Hymenobacter*, *Sphingomonas*, *Bdellovibrio*, and *Terriglobus*, were detected in 100% of the samples (Fig 5D). These core taxa are likely important contributors to host health, potentially involved in nutrient cycling, stress tolerance, and protection against pathogens. Their consistent presence across different plant genera suggests a key role in maintaining the stability and resilience of the Myrtaceae-associated microbial community.

Additionally, approximately 0.73% of the bacterial sequences remained unclassified at the phylum level, indicating the presence of potentially novel or poorly characterized taxa within these plant-associated communities.

Functional prediction of the bacterial community associated with Myrtaceae plants revealed that the predominant predicted functions included aerobic and general chemoheterotrophy, fermentation, hydrocarbon degradation, methanotrophy, methylotrophy, nitrogen fixation, and ureolysis, all of which are potentially important for plant growth and ecosystem functioning (Fig S3; Table S1).

## 4. Discussion

Plants produce chemical signals to attract specific microbiota to their tissues, which in turn provide protection against herbivory and pathogens, while also enhancing various physiological traits (reviewed in Pang et al., 2021)^23^. Here, we provide a comprehensive characterization of the bacterial communities inhabiting the phyllosphere of *Eugenia uniflora* (pitanga) and its Myrtaceae relatives. Given the pharmacological and ecological importance of Myrtaceae, understanding their microbiomes is particularly relevant for both basic and applied research ^24,25^.

Previous microbiome studies in Myrtaceae have focused largely on fungi. For example, fungal endophytes have been described in *Metrosideros excelsa* ^26^, *Eucalyptus* ^27–29^, and *Eugenia* species ^30^, with an emphasis on culturable taxa. Similarly, the fungal community of *Myrtus communis* has been characterized using high-throughput sequencing ^31^. By contrast, bacterial phyllosphere communities in Myrtaceae remain poorly documented, with most efforts in the Atlantic Forest restricted to culturable bacteria from *Eucalyptus*, *Eugenia*, and *Plinia* ^13,32,33^. Our study therefore fills an important gap by applying high-throughput sequencing to describe both culturable and unculturable bacteria across developmental stages of *E. uniflora* and in some Myrtaceae genera. This approach is significant from ecological, agronomical, and biotechnological perspectives, providing valuable insights into the microorganisms coexisting with this group of plants.

### 4.1 Pitanga developmental stage shapes bacterial community structure and function

The phyllosphere of mature *E. uniflora* trees exhibited greater alpha diversity compared to young plants, and developmental stage significantly shaped beta diversity, indicating distinct microbial assemblies. This reinforces the idea that plant ontogeny is a key determinant of microbiome composition ^34–36^.

The observed differences likely arise from age-dependent physiological and biochemical factors. Mature leaves generally present larger surfaces, longer exposure to environmental microbial pools, and higher accumulation of organic compounds and exudates, creating a more heterogeneous and microbially favorable environment ^37^. Consistently, our functional predictions indicated that microbes from mature leaves were enriched in pathways associated with secondary metabolite metabolism, whereas those from young leaves were characterized by pathways related to microbial communication and defense. This suggests that shifts in chemical composition during plant development strongly influence microbial recruitment and functional potential ^38^.

From an ecological perspective, the higher microbial diversity of mature trees may provide increased functional redundancy and resilience, improving nutrient acquisition, disease resistance, and stress tolerance ^39^. Together, these findings highlight plant developmental stage as a fundamental driver of microbiome structure and function, reinforcing the holobiont concept and its importance in ecological and agricultural contexts.

### 4.2 A core microbiome underpins pitanga and its Myrtaceae relatives

Across *Eugenia uniflora* and other Myrtaceae, we identified a conserved core microbiome dominated by Proteobacteria, Bacteroidetes, and Actinobacteriota, which together accounted for over 95% of sequencing reads. At the genus level, *Methylobacterium–Methylorubrum*, *Hymenobacter*, *Sphingomonas*, *Bdellovibrio*, and *Terriglobus* were consistently detected across all samples. The persistence of these taxa across different hosts suggests that coevolutionary processes and adaptation to shared leaf-surface conditions, such as UV exposure, desiccation, and interactions with secondary metabolites, play a key role in shaping the Myrtaceae phyllosphere ^40^.

*Methylobacterium–Methylorubrum*, well-known phyllosphere colonizers, utilize single-carbon compounds released during plant growth and senescence, promoting plant growth and enhancing tolerance to abiotic stress ^41–43^. Similarly, *Hymenobacter* and *Sphingomonas* contribute to plant health through stress mitigation and by modulating bioactive compound production, which may influence plant defense and physiology ^44–46^. *Bdellovibrio*, a predatory bacterium, can regulate microbial community composition by preying on Gram-negative bacteria, potentially suppressing pathogens and maintaining microbiome stability ^47,48^. *Terriglobus* contributes to carbon cycling, nutrient mobilization, and soil stability, acting as a metabolic backbone that supports both rhizosphere and phyllosphere function ^49–51^.

The consistent occurrence of these genera across Myrtaceae highlights their potential roles in host adaptation, resilience, and overall ecosystem functioning. These taxa may represent promising candidates for biotechnological applications, including biocontrol, biofertilization, and phytoremediation strategies.

However, it is important to note that the functional roles inferred from literature and sequence-based predictions require experimental validation in *E. uniflora* specifically. Core microbiome identification can also be influenced by sampling strategy, sequencing depth, and primer choice, which should be considered when interpreting these patterns. Building on these considerations, our study has additional limitations. First, by relying solely on 16S rRNA amplicon sequencing, we achieved genus-level resolution but could not fully capture strain-level diversity or directly confirm functional predictions. Second, sampling was restricted to a single site in the Atlantic Forest biome, which may not reflect broader environmental or geographic variation in Myrtaceae microbiomes. Third, we focused on two developmental stages as discrete time points, without assessing seasonal or longitudinal dynamics. Finally, while we targeted the phyllosphere (epiphytic and endophytic compartments), belowground compartments such as the rhizosphere were not included, even though they may strongly interact with foliar microbiota. Addressing these limitations through shotgun metagenomics, metabolomics, multi-site sampling, and integrative plant–microbe studies will be crucial to refine our understanding of Myrtaceae-associated microbiomes and validate the ecological roles of the conserved core taxa identified here.

## 5. Conclusions and perspectives

This study provides the first high-throughput sequencing–based profile of the Myrtaceae phyllosphere microbiome, demonstrating that developmental stage is a major determinant of bacterial diversity and function in *E. uniflora.* The identification of a conserved core microbiome across Myrtaceae points to long-term plant–microbe associations that may underpin host health and stress tolerance. Beyond ecological insights, these findings emphasize the potential of Myrtaceae as reservoirs of beneficial bacteria with applications in bioinoculant development, natural product discovery, and sustainable agriculture. Future research combining metagenomics, metabolomics, and functional validation will be crucial to uncover the mechanistic basis of these associations and translate them into applied outcomes.

## ACKNOWLEDGMENTS

Financial support was provided by the Universidad Peruana de Ciencias Aplicadas (internal fund A-065-2024 to FG) and INCT-MCTI/CNPq/CAPES/FAPs n°16/2014 for the conduct of the research and preparation of the article.

## DATA AVAILABILITY

The raw sequence data are publicly available in the GenBank database under BioProject accession number PRJNA1122548.

## AUTHOR CONTRIBUTIONS

**ICCS**: Conception and design, acquisition of data, application of computational techniques to analyze the data, interpretation of results and paper writing. **DEB**: Application of computational techniques to analyze the data and interpretation of results. **RM**: Conception and design, acquisition of financial support, project supervision and interpretation of results. **FLGE:** Conception and design, acquisition of financial support, application of computational techniques to analyze the data, project supervision and interpretation of results. All the authors read and approved the manuscript.

## STATEMENT

During the preparation of this work the author(s) used ChatGPT in order to check grammar. After using this tool/service, the author(s) reviewed and edited the content as needed and take(s) full responsibility for the content of the publication.

## Supplementary material

**Fig S1**. **Taxonomic composition of the *Eugenia uniflora* phyllosphere bacterial community.** (a) Relative abundance of major bacterial phyla showing dominance of *Proteobacteria*, *Bacteroidota*, and *Actinobacteriota*. (b) Taxonomic distribution at the family level.

**Fig S2. Taxonomic composition of the Myrtaceae phyllosphere bacterial community.**

(a) Relative abundance of major bacterial phyla showing dominance of *Proteobacteria*, *Bacteroidota*, and *Actinobacteriota*. (b) Taxonomic distribution at the family level.

**Fig S3. Functional profile of the Myrtaceae phyllosphere bacterial community based on FAPROTAX analysis.** Bar plots represent the relative abundance of predicted ecological functions across plant samples.

## List of Supplementary Table

**Table S1.** ASV abundances per sample from *Eugenia uniflora*.

**Table S2.** ASV abundance per sample of Myrtaceae plants.

**Table S3.** Alpha and beta diversity statistical analysis of *Eugenia uniflora* plants.

**Table S4.** Bacterial abundance at the genus level in *Eugenia uniflora* and other Myrtaceae plants.

**Table S5.** Bacterial genera unique or shared across plant ages based on Venn diagram analysis.

**Table S6.** Bacterial genera with differential abundance across *Eugenia uniflora* ages identified by edgeR.

**Table S7.** Metabolic functions across *Eugenia uniflora* and other Myrtaceae plants predicted by Tax4Fun.

**Table S8.** Ecological functions across Eugenia uniflora plant ages predicted by FAPROTAX.

**Table S9.** Alpha and beta diversity statistical analysis for Myrtaceae plants.

**Table S10.** Bacterial occupancy and relative abundance in the Eugenia uniflora microbiome.

**Table S11.** Bacteria occupancy and relative abundance from Myrtaceae microbiome.

**Table S12.** Ecological functions across Myrtaceae predicted by FAPROTAX.

